# A single amino acid change drives left- right asymmetry in Bilateria

**DOI:** 10.1101/2022.10.06.511172

**Authors:** Marta Truchado-García, Kimberly J. Perry, Florencia Cavodeassi, Nathan J. Kenny, Jonathan Q. Henry, Cristina Grande

**Affiliations:** Departamento de Biología, Facultad de Ciencias, Universidad Autónoma de Madrid, Madrid, Spain; University of California – Berkeley, Department of Molecular and Cell Biology, 571 Weill Hall, Berkeley, CA 94702, USA; University of Illinois, Department of Cell and Developmental Biology, 601 S. Goodwin Avenue, Urbana, IL 61801, USA; Institute for Medical and Biomedical Education, St. George’s University of London, Cranmer Terrace, London, SW17 0RE, UK; Natural History Museum, Cromwell Road, London, UK; Department of Biochemistry, University of Otago, P.O. Box 56, Dunedin, Aotearoa-New Zealand; The Marine Biological Laboratory, University of Chicago, 7 MBL St, Woods Hole, MA 02543, USA

**Keywords:** Nodal, EGF-CFC, Cripto, left-right asymmetry, Zebrafish, Evo-Devo, gene expression pattern, Spiralia, *Crepidula fornicata*

## Abstract

Asymmetries are essential for proper organization and function of organ systems. Genetic studies in deuterostomes have shown signaling through the Nodal/Smad2 pathway plays a key, conserved role in the establishment of body asymmetries. While Nodal signaling is required for the establishment of left-right asymmetry (LRA) across bilaterian species, little is known about the regulation of Nodal signaling in spiralians. Here, we identified orthologs of the *egf-cfc* gene, a master regulator of the Nodal pathway in vertebrates, in several invertebrate species, the first evidence of its presence in non-deuterostomes. Our functional experiments indicate that despite being present, *egf-cfc* does not play a role in the establishment LRA in gastropods. However, experiments in zebrafish suggest that a single amino acid mutation in the *egf-cfc* gene in the deuterostome common ancestor was the necessary step in inducing a gain-of-function in LRA regulation. This study shows that that the *egf-cfc* gene likely appeared in the bilaterian stem lineage, before being adopted as a master mechanism to regulate the Nodal pathway and the establishment of LRA in deuterostomes.

## INTRODUCTION

Despite dramatically different body architectures, animals share common signaling pathways and transcriptional networks that regulate their development, a core “genetic toolkit” (Cheatle-Jarvela & Pick, 2016). Here we examine the genetic toolkit that underlies left-right asymmetry (LRA) in the Spiralia, a group that contains molluscs, annelids, platyhelminthes, and at least 8 other phyla, and is the most phyla-rich and arguably morphologically diverse bilaterian clade (Marlétaz *et al*., 2019).

Most bilaterians exhibit some left-right asymmetries in both their external and internal anatomy (e.g., the position of the heart and liver in vertebrates, and the direction of gut torsion in vertebrates and arthropods). Both paired and unpaired organs generate asymmetries, according to their shape, configuration and function. These asymmetries are essential for proper packing connectivity and function of these organ systems. Complete reversal of these asymmetries (i.e., *situs inversus*) to form a perfect mirror image is often well tolerated, but partial reversals (e.g. heterotaxies) may severely compromise viability (Lobikin *et al*., 2012). Within the Spiralia, shell coiling of gastropod snails is perhaps the most obvious morphological asymmetry, but others include the positions of certain sensory organs, various neural connections, the digestive tract and locomotive cilia (Grande, 2010; Martín-Durán *et al*., 2016). Most snails present a coiled shell (and by extension, body conformation) that has a clear chirality – either dextral (with a clockwise twist, as is more common) or sinistral (with an anti-clockwise twist). However, chirality is evident during embryonic development, during the second and third cleavage divisions; and these define the chirality of the embryo and its future larval/adult body plan (Shibazaki *et al*., 2004). In gastropods, a dextral third cleavage results in animals with dextral shells and vice versa.

The establishment of LRA has been studied mainly in vertebrate models and in tissue culture, although there are several studies in invertebrates (reviewed in Hamada & Tam, 2020). These studies have demonstrated that symmetry breaking starts very early in embryogenesis. The information generated by the early symmetry breaking events is translated into asymmetric deployment of genetic cascades, reinforcing the initial signal, and reliably dictating subsequent asymmetries during organogenesis. At the molecular level, the initial determination of LRA in vertebrates is widely conserved. However, conservation and divergences in these mechanisms among metazoans requires further analysis (Palmquist & Davidson, 2017). Signaling through the Nodal/Smad2 pathway plays a role in LRA in diverse deuterostomes (Duboc *et al*., 2005; Soukup *et al*., 2015) and some spiralians (Grande & Patel, 2009; Grande *et al*., 2014; Martín-Durán *et al*., 2016). This pathway controls cellular processes such as differential cell migration, proliferation, adhesion and asymmetric morphogenesis in both deuterostomes and the Spiralia (Hamada *et al*., 2002; Duboc *et al*., 2005: Grande & Patel, 2009). Indeed, loss of function (LOF) of components in the pathway can promote *situs inversus* or heterotaxy (Levin *et al*., 1997; Grande & Patel, 2009). However, the Nodal network is absent in Ecdysozoa, which suggests independent evolution of the mechanisms driving LRA in this clade (Schier, 2009).

*nodal* belongs to the TGFβ superfamily of signalling molecules, and signals asymmetrically along a single side of the developing embryo, through a well-conserved cascade (Massagué, 2012), to create a phospho-Smad complex, which promotes the activation of target genes, such as *Pitx*. Unlike in other TGFβ pathways, an EGF-CFC (Epidermal Growth Factor-CFC) coreceptor is required for the assembly of the Nodal receptor complex. Comparative studies have demonstrated similar expression and regulatory interactions between *nodal* and *Pitx* in diverse deuterostomes (reviewed in Swalla, 2006; Palmquist & Davidson, 2017). In snails, chirality defines the location of Nodal activation, so that the gene is expressed on the right side of dextral and the left side in sinistral individuals (Grande & Patel, 2009; Kuroda *et al*., 2009). Defective function of the Nodal pathway leads to LRA defects in snails (Grande & Patel, 2009). However, little is known about the regulation of Nodal signalling in spiralians.

EGF-CFC coreceptors play a key role in the regulation of Nodal activity. These signalling factors have diverse functions, from controlling early events during embryonic development to tissue homeostasis. The EGF-CFC protein contains recognition sites for interactions with diverse molecules. For instance, they are able to block signaling of TGF-β ligands such as Activin or enhance the signaling of other ligands such as BMP-4 or Nodal (Gray & Vale, 2010; Minchiotti *et al*., 2001; Yan *et al*., 2002). Three main domains define this protein family (fig. 3B): 1) The EGF-like domain, involved in ligand recognition (i.e TGF-βs, BMPs, GDFs and Tomoregulin), and regulation of the Nodal, MAPK and PI3K-Akt pathways; 2) the CFC domain, involved in receptor binding; and 3) the GPI domain, involved in membrane anchoring (reviewed in Klauzinska *et al*., 2014). To date, EGF-CFCs have only been described in deuterostomes, and are considered an evolutionary innovation of the group (reviewed in Shen & Schier, 2000; Ravisankar *et al*., 2011). In mammals, the family includes two genes called *cripto* (Ding *et al*., 1998), and *cryptic* (Yan *et al*., 1999). In the zebrafish (*Danio rerio*), one EGF-CFC gene has been described (*one-eye-pinhead, oep*). *Oep* is maternally and zygotically expressed; at later stages it is expressed in the forebrain, lateral plate and notochord. Zygotic *oep* mutants show many of the observed phenotypes for *cripto* and *cryptic* mutants in mice, including defects in the establishment of both left-right and anterior-posterior axes (Schier *et al*., 1997).

The establishment of LRA is one of the most studied functions of EGF-CFC (reviewed in Klauzinska *et al*., 2014). The interaction between EGF-CFC and Nodal at a molecular level is well described in both mice and humans (Calvanese *et al*., 2010). A threonine within the EGF-like domain (position T88 in humans and T72 in mouse) is essential for Nodal function (fig. 3C). This residue undergoes O-fucosilation, facilitating the ligand-coreceptor interaction. This O-fucosilation cannot be replaced by any other amino acid, even those capable of being fucosilated (serine) (Schiffer *et al*., 2001; Shi *et al*., 2007). Whether this interaction is also necessary for the activation of the Nodal pathway in other organisms remains unknown.

We have surveyed the presence and/or role in LRA of both Nodal and the EGF-CFC coreceptor in non-deuterostomes, focusing on gastropods, to understand the relationship between these two proteins in non-deuterostomes. We identified orthologs of EGF-CFC in several invertebrate species, the first evidence of its presence in non-deuterostomes. We infer that, despite *egf-cfc* orthologs being present in non-deuterostomes, the Nodal pathway may have acted in an *egf-cfc*-independent manner in the common ancestor of all bilaterians, and this relationship persists in non-deuterostomes. Our experiments in Zebrafish suggest that the role of Nodal in LRA asymmetry is shared between deuterostomes and non-deuterostomes, although the pathway became dependent on the EGF-CFC coreceptor in the deuterostome ancestor sometime after a single key amino acid change occurred in its protein sequence.

## RESULTS

### Two paralogs for *nodal* and one ortholog for *egf-cfc* are present in *C. fornicata*

Searches in the gastropod *C. fornicata* transcriptome database and subsequent orthology analyses identified two paralog genes for *nodal* (*Cfo-nodalA* and *Cfo-nodalB*) (figs. 1A and 1B) (Grande *et al*., 2014). *Cfo-nodalA* gives rise to a 428 amino acid protein, while *Cfo-nodalB* generates a 472 amino acid protein. Both proteins display the distinctive seven cysteine residues that exist in the C-terminal region of this family of proteins, and the cleavage site domain, which matches orthologs in other animals. We observe a similar duplication in another gastropod, *Patella vulgata*, but the phylogenetic analysis performed in Grande *et al*. (2014) suggests these paralogs may have arisen independently.

**Figure 1.**
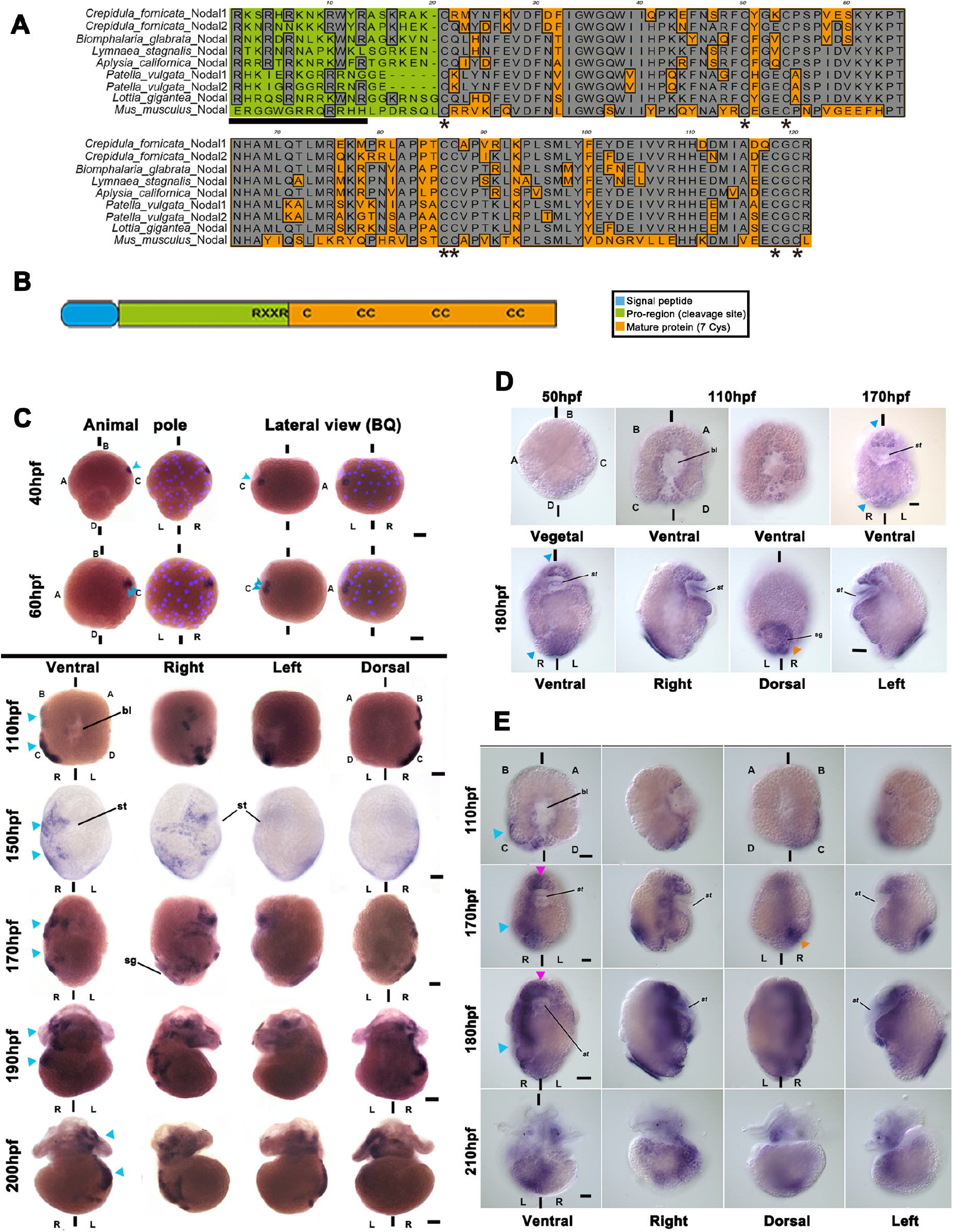
Two paralogs for *nodal* are present in *C. fornicata*, and *Cfo-nodalB* expression suggests involvement in left/right asymmetry development. A) Alignment of several ortholog proteins for Nodal in different gastropod species (*Crepidula fornicata; Biomphalaria glabrata; Lottia gigantea; Patella vulgata; Aplysia californica; Littorina stagnalis*) and mammalian *Mus musculus*. Grey indicates conservation for one given position; asterisks point the seven conserved cysteines that characterize the family; black bar delimits the regions were cleavage sites are located. B) Generic structure of Nodal protein. (RXXR): cleavage site; C: cysteines. C-E) Spatio-temporal localization of (C) *Cfo-nodalB*, (D) *Cfo-nodalA*, and (E) *Pitx* mRNA by whole mount *in situ* hybridization during the development of *C. fornicata*. Several views of the same embryo for each stage are indicated on the figure. st, stomodeum; bl, blastopore; sg, shell gland; L, left; R, right. A, B, C and D indicate the quadrants; hpf, hours post fertilization; bilateral axis is represented by the vertical bars. Scale corresponds to 30 μm. In C) Superpositions of brightfield images with Hoechst staining have been included for early stages, to identify the cells. Note that the expression is asymmetrical in all the stages (blue arrowheads). D) Note that the expression is symmetrical in all the stages (blue arrowheads). orange arrowhead, shell gland. E) Pink arrowhead, symmetrical signals; blue arrowhead, asymmetrical signals; orange arrowhead, shell gland.

Next, we focused on the EGF-CFC family (reviewed in Gray & Vale, 2012). Comprehensive searches in metazoan databases for the *egf-cfc* orthologs and subsequent orthology analyses were carried out using *cripto* and *cryptic* genes from *Mus musculus*, and *oep* from *D. rerio* as queries. We found orthologous genes for the *egf-cfc* gene in gastropod molluscs, annelids and brachiopods (figs. 3A and 3C; Supp. material S2), while exhaustive searches of cnidarian and ctenophore genomes, genomes of bilaterians such as *Xenoturbella*, Acoelomorpha, Ecdysozoa and Platyhelminthes, and the NCBI nr, TSA and EST databases, retrieved no orthologs as members of this family. This result indicates that the last ancestor of Bilaterians presented only one EGF-CFC member, lost in the Ecdysozoan lineage, and that the EGF-CFC family is not an evolutionary novelty of deuterostomes. While only one representative *egf-cfc* gene was found in most bilaterians, our phylogenetic analysis shows the existence of up to 3 paralogous genes for *egf-cfc* within specific evolutionary lines (*Xenopus laevis, X. tropicalis, Homo sapiens* and *P. vulgata*) (fig. 3A). The phylogenetic analysis also suggests a duplication in the deuterostome lineage, which gave rise to another clade of proteins in mammals (Cripto).

Sequence analysis showed that all orthologs display a conserved structure that includes an EGF-like domain, a CFC domain and a GPI domain (figs. 3B and 3C), generating a 209 amino acid protein in *C. fornicata* that we call Cfo-EGF_CFC. Three critical amino acids in the EGF-like domain of the EFG-CFC coreceptor (Bianco *et al*., 2002; Yan *et al*., 2002; Yeo & Withman, 2001) have been identified in humans for correct interaction with Nodal: G87, T88 and F94 (Minchiotti *et al*., 2001; Shaolin *et al*., 2007). The substitution of these amino acids leads to a complete loss of function of EGF-CFC. Three-dimensional models have confirmed that these three amino acids are exposed on the same side of the protein and form a surface that may serve as an interaction point with Nodal. G87 and F94 are conserved in all the orthologs identified here (fig. 3C). However, T88 is only conserved in deuterostomes, whereas gastropod mollusks have a leucine (L) or arginine (R) at this position, annelids have a methionine (M) or isoleucine (I), and brachiopods have a valine (V) (Fig. 3C). Previous work (Schiffer *et al*., 2001; Yan *et al*., 2002; Shaolin *et al*., 2007) point at T88 as being critical for Nodal-Cripto interaction, and its substitution by other amino acids leads to a failure on the activation of the pathway.

The CFC domain mediates the protein-protein interaction of EGF-CFC with the type-I receptor of the TGF-β pathway (Alk4) (Calvanese *et al*., 2009), in particular, positions H120 and W123 in humans (Foley *et al*., 2003; Marasco *et al*., 2006). Our analysis shows that these are conserved in all the identified orthologs, with the exception of W123 for the annelid *Spirobranchus*, and hence, all of them presumably bind to the type-I receptor (fig. 3C).

Last, these proteins can act not only as membrane proteins but also as soluble proteins, if a cleavage on the GPI domain occurs as shown in human cell cultures (Watanabe & Salomon, 2010). Our results showed the existence of a GPI domain in all the identified orthologs, so these orthologs may share the ability to act as either membrane or soluble proteins.

### *Cfo-nodalB* expression suggests an involvement in LRA development

We designed specific riboprobes for *Cfo-nodalA* and *Cfo-nodalB* by selecting regions within the sequences that were divergent enough to avoid a cross-reaction. Whole mount *in situ* hybridization (WISH) was subsequently carried out at different stages in *C. fornicata* embryos to examine their expression throughout development. *Cfo-nodalA* transcript is clearly present at the mid-gastrula stage (110 hours post fertilization; hpf) in every quadrant (A, B, C and D) and around the blastopore (fig.1D), which will form the mouth in these animals. At late gastrula stages, when organogenesis starts (170 hpf), *Cfo-nodalA* is located along the anterior end and around the stomodeum (future mouth), and in a small posterior region (fig. 1D, blue arrowhead). In the pre-veliger larva (180 hpf), *Cfo-nodalA* expression is more intense and remains restricted to an area of the mouth in a bilaterally symmetrical manner, as well as in the shell gland (fig.1D; orange arrowhead). *Cfo-nodalB* is detected when embryos reach 40-50 cells (40 hpf) (fig. 1C; blue arrowhead). At this stage expression is detected in a single cell of the C quadrant (2c^2^). As subsequent divisions take place, *Cfo-nodal2* appears in the 2c^2^ cell progeny (2c^21^ and 2c^22^), at 60 hpf (fig. 1C; blue arrowhead). During gastrulation (110 hpf), the expression remains in the progeny of these cells (fig. 1C). During organogenesis and larval development (from 150 hpf), this expression is asymmetrical, located on the right side of the mouth, the right area of the velum, the rudiment of the foot and in an ectodermal patch on the right side of the post-trochal region (fig. 1C; blue arrowhead). This expression becomes more intense and remains in these regions of pre-veliger stages (fig. 1C). Finally, in veliger larvae (200 hpf), expression is restricted to the right side of the visceral mass and the head (fig. 1C). Thus, while *Cfo-nodalA* expression is largely symmetrical, *Cfo-nodalB* shows clear asymmetric expression, concentrated on the right side of the developing embryo (figs. 1D and 1C).

*Cfo-nodalB* expression in the right side of the embryo is closely followed by asymmetric expression of *Pitx*, a known target gene of Nodal in gastropods and other bilaterians (Grande & Patel, 2009; Hamada *et al*., 2002). *Pitx* is asymmetrically expressed in *C. fornicata* immediately after *Cfo-nodalB* activation (fig. 1E), which is consistent with previous data in other gastropods (Grande & Patel, 2009). *Pitx* expression is asymmetrical on the right side as previously shown in Henry *et al*., 2010. This expression on the right side is evident at mid-gastrula stages (fig. 1E; blue arrowhead), and colocalizes to the same regions of *Cfo-nodalB* (figs. 1C and 1E). *Pitx* shows two expression sets, one asymmetrical on the right side, and another symmetrical, as is the case in other gastropods and vertebrates, suggesting the presence of independent enhancer areas that regulate its expression (Christiaen *et al*., 2005; Grande & Patel, 2009).

### *Cfo-nodalB* regulates the establishment of LRA and Activin inhibits the pathway in *C. fornicata*

To further investigate the function of *Cfo-nodalB*, we produced wild-type and defective forms for expression in *C. fornicata*. Overexpression experiments carried out through microinjection of an *in vitro* synthesized mRNA for the *Cfo-nodalB* paralog into *C. fornicata* zygotes (table 1) produced a high percentage of affected individuals. We observed a consistent phenotype, including loss of LRA in the body plan. This included a symmetrical shell and arrangement of the internal organs (i.e., no hindgut rotation with the lack of torsion) (fig. 2B; table 1). The scoring of the embryos with a symmetrical phenotype was dose-dependent and directly related to the amount of RNA injected into the embryos. At the highest injection concentration, 92% of embryos were symmetrical (table 1).

**Table 1.**
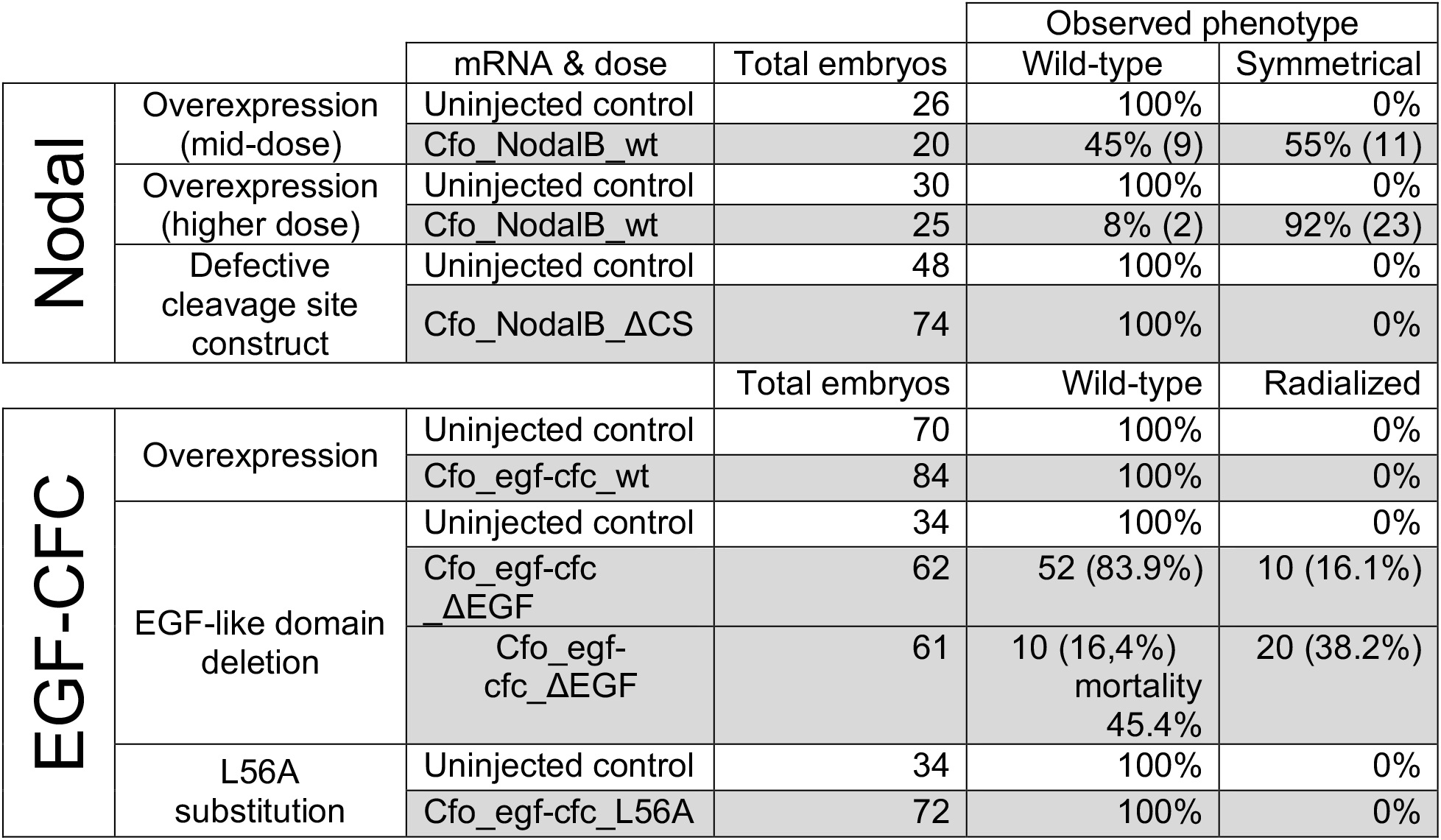
Results obtained after mRNA injection on *C. fornicata* zygotes, of *nodalB* and *egf-cfc* constructs.

**Figure 2.**
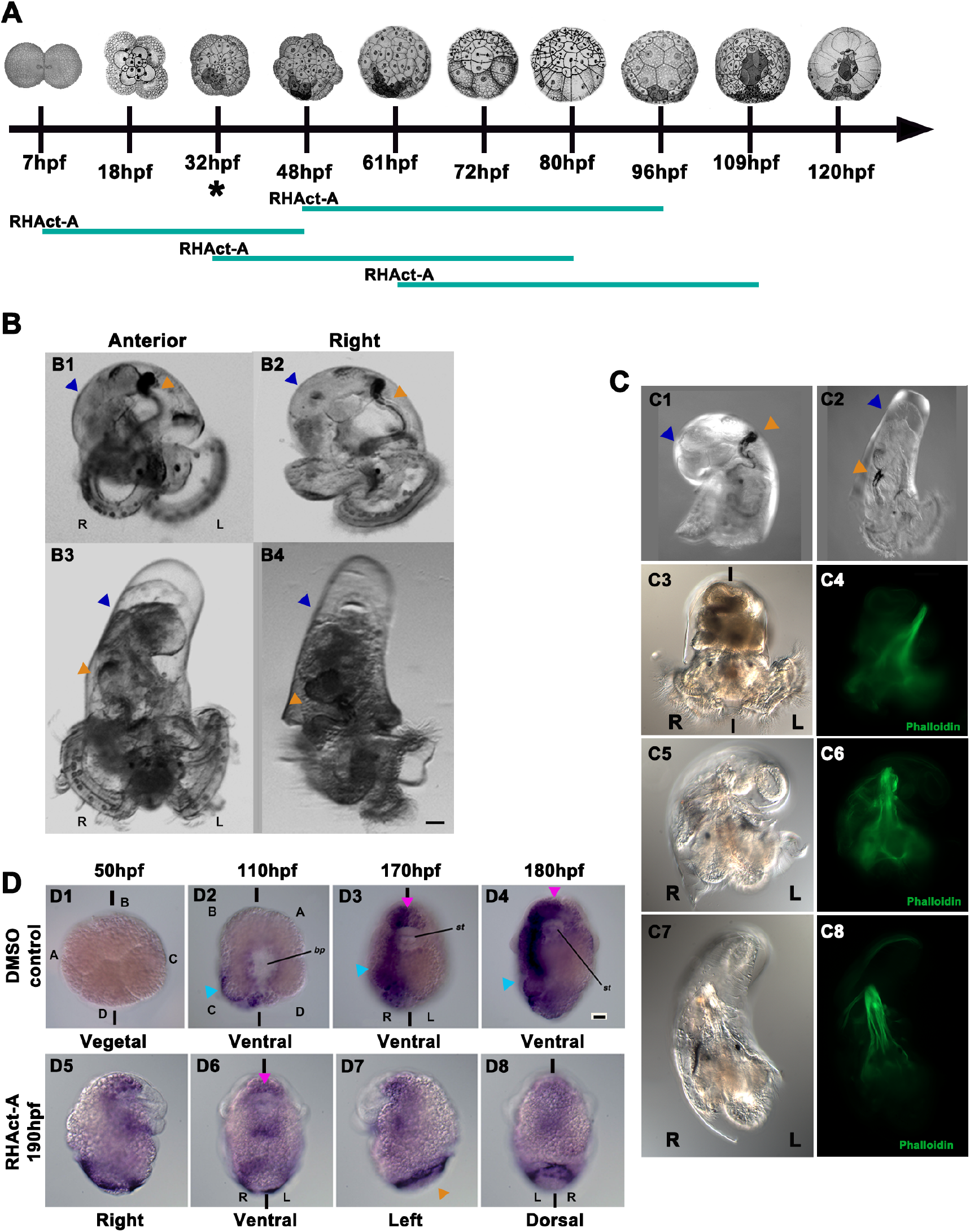
*Cfo-nodalB* regulates the establishment of LRA, while Activin inhibits the Nodal pathway in *C. fornicata*. A) Diagram of the windows used on RHAct-A treatment. The teal bar represents the time in which the embryos were cultured in RHAct-A, and the asterisk indicates when NodalB expression first starts; hpf, hours post-fertilization. B) Overexpression of *Cfo-nodalB*, in a 2-weeks old hatched veliger. (B1) Anterior and (B2) right views of a control veliger; (B3) anterior and (B4) right views of a veliger after *Cfo-nodalB* injection. C) RHAct-A treatment, in a 2-weeks old hatched veliger. Right view of a (C1) control veliger and (C2) a treated veliger. C3-C8) Three individuals with different grades of coiling after treatment. Bright-field of (C3) coiled DMSO control, (C5) hemitorsioned and (C7) straight larvae after RHAct-A. C4/C6/C8: Note the change of the retractor muscle of those same larvae (C3, C5, C7) shown by phalloidin stain. In both B and C, Note the bilateral symmetry phenotype in B3, B4 and C2, C7. L, left; R, right; shell coiling, blue arrowhead; gut coiling, orange arrowhead. D) Spatio-temporal localization of *Pitx* mRNA by WISH, in RHAct-A treated *C. fornicata* embryos. D1-D4) Control embryos in vegetal/ventral view in four stages during development. D1, non-stained blastula (50 hpf); D2, mid-gastrula (110 hpf); D3, organogenesis (170 hpf); D4, preveliger larva (180 hpf). Note the right-side staining (cyan arrowhead) and the symmetrical staining (pink arrowhead) on the anterior region and stomodeum (D2-D4). D5-D8) Views of a preveliger, post-RHAct-A exposure. Note that the symmetrical expression remains (pink arrowhead), while the asymmetrical expression is lost on the right side; shell gland, orange arrowhead.st, stomodeum; bl, blastopore; L, left; R, right. A, B, C and D indicate the quadrants; bilateral axis is represented by the vertical bars. Scale bars correspond to 30 μm.

Next, we investigated whether the removal of the putative cleavage sites within the *Cfo-nodalB* precursor would inhibit activation of the *Cfo-nodalB* pathway (fig. 1B). The cleavage and dimerization of the ligands are necessary for these to bind to the corresponding receptors (Shen, 2007). Such changes are effective in loss-of-function experiments, since their presence leads to competition with wild-type forms that block the activation of functional dimers (Hawley *et al*., 1995; Joseph & Melton, 1998). This potentially dominant negative form was designed to disrupt Nodal signaling at the ligand-receptor level. However, when this modified mRNA was injected in *C. fornicata* zygotes, no phenotypes different from wild-type were observed in resulting larvae at any concentration (table 1). The utility of this approach may be questioned by studies performed in *Xenopus*, which showed that even non-processed peptides can generate some signaling (Eimon & Harland, 2002). However, our result suggests that our design may result in a non-functional product, given the lack of an overexpression phenotype.

As a second approach to manipulate the Nodal pathway, we treated the embryos with recombinant Activin. Activin is a TGF-β ligand closely related to Nodal and is able to functionally replace it *in vitro* (Namwanje & Brown, 2016), but interestingly, it is not present in gastropods (Grande & Patel, 2009; Grande *et al*., 2014). To date, no ortholog of Activin has been identified for Spiralia, but there are several Activin-like paralogs that result from duplication and divergence events of the ancestral gene (Kenny *et al*., 2014). Some of these paralogs have been analyzed and it has been demonstrated that these are not active during gastropod development (Grande & Patel, 2009). Hence, the available evidence suggests the absence of a role in gastropod development for these Activin paralogs. Under these conditions, it is expected that the recombinant human Activin A (RHAct-A) may compete with Nodal for receptors in *C. fornicata* embryos without promoting signal transduction (Supp. material S1)

RHAct-A treatments administered to 2-cell stage embryos (7hpf) to 48hpf did not show symmetrical larvae. However, embryos treated at 32 hpf (the stage of *Cfo-nodalB* activation in *C. fornicata* according to our WISH results) presented the symmetrical body plan observed in the overexpression experiment (symmetrical shells and loss of torsion in the visceral mass), in a dose-dependent manner in up to 80% of the cases at the highest dose (figs. 2A and 2C; table 2). Thus, the RHAct-A treatment during this window of time leads to a symmetric body plan, presumably due to RHAct-A outcompeting the endogenous Nodal ligands (Supp. material S1). On the other hand, when we shifted the window to 48 hpf and 61 hpf, this percentage decreased to around 60% of cases. The lack of an observable phenotype at 7 hpf treatment can be explained by the fact that *Cfo-nodalB* expression is not present until at least until 40 hpf, only 8 hours prior to removing the treatment (fig. 1C). Additionally, the only cell expressing the ligand at that time is in fact producing Nodal, and the results suggest that the early removal of the RHAct-A at that time compensates for the competition with the beginning of *Cfo-nodalB* expression, which eventually reaches its physiological concentration.

**Table 2.** Results obtained after RHAct-A treatment on *C. fornicata* embryos.

| Window | Concentration (μM) | In | Out | Total embryos | Observed phenotype |  |
| --- | --- | --- | --- | --- | --- | --- |
|  |  |  |  |  | Wild-type | Symmetrical |
| Cleavage (7 hpf) | Control- 0 | 7h | 48h | 65 | 65 | 0 |
|  | 10 |  |  | 56 | 56 | 0 |
| Early blastula (32 hpf) | Control- 0 | 32h | 80h | 65 | 65 | 0 |
|  | 10 |  |  | 51 | 10 | 41 (80.4%) |
| Blastula (48 hpf) | Control- 0 | 48h | 96h | 70 | 70 | 0 |
|  | 5 |  |  | 45 | 41 | 4 (8.8%) |
|  | 10 |  |  | 43 | 18 | 25 (60%) |
| Gastrula (61 hpf) | Control- 0 | 61h | 109h | 30 | 0 | 0 |
|  | 10 |  |  | 32 | 12 | 20 (62.5%) |

Analysis of *Pitx* expression revealed that RHAct-A was producing an inhibition phenotype, instead of an overexpression effect (fig. 2D). Indeed, although the symmetrical *Pitx* domain in the anterior region persists unaffected, the asymmetric domain is lost in those larvae treated with RHAct-A (fig. 2D). Additionally, the retractor muscle (fig. 2C3-2C8) is straight in comparison with control larvae, suggesting a lack of torsion. Thus, the presence of the RHAct-A protein interferes with LRA in the gastropod *C. fornicata*, when applied after the 32 hpf stage, potentially acting as a dominant-negative inhibitor that interferes with Nodal signaling (Supp. material S1). *Cfo-nodalB* expression continues until later stages (200 hpf), colocalizing with the *Pitx* signal (figs. 1C and D). However, the decrease of symmetrical individuals in blastula treatments suggests that the initial activation of the pathway between 32 hpf and 61 hpf may be key for the correct establishment of LRA.

**Figure 3.**
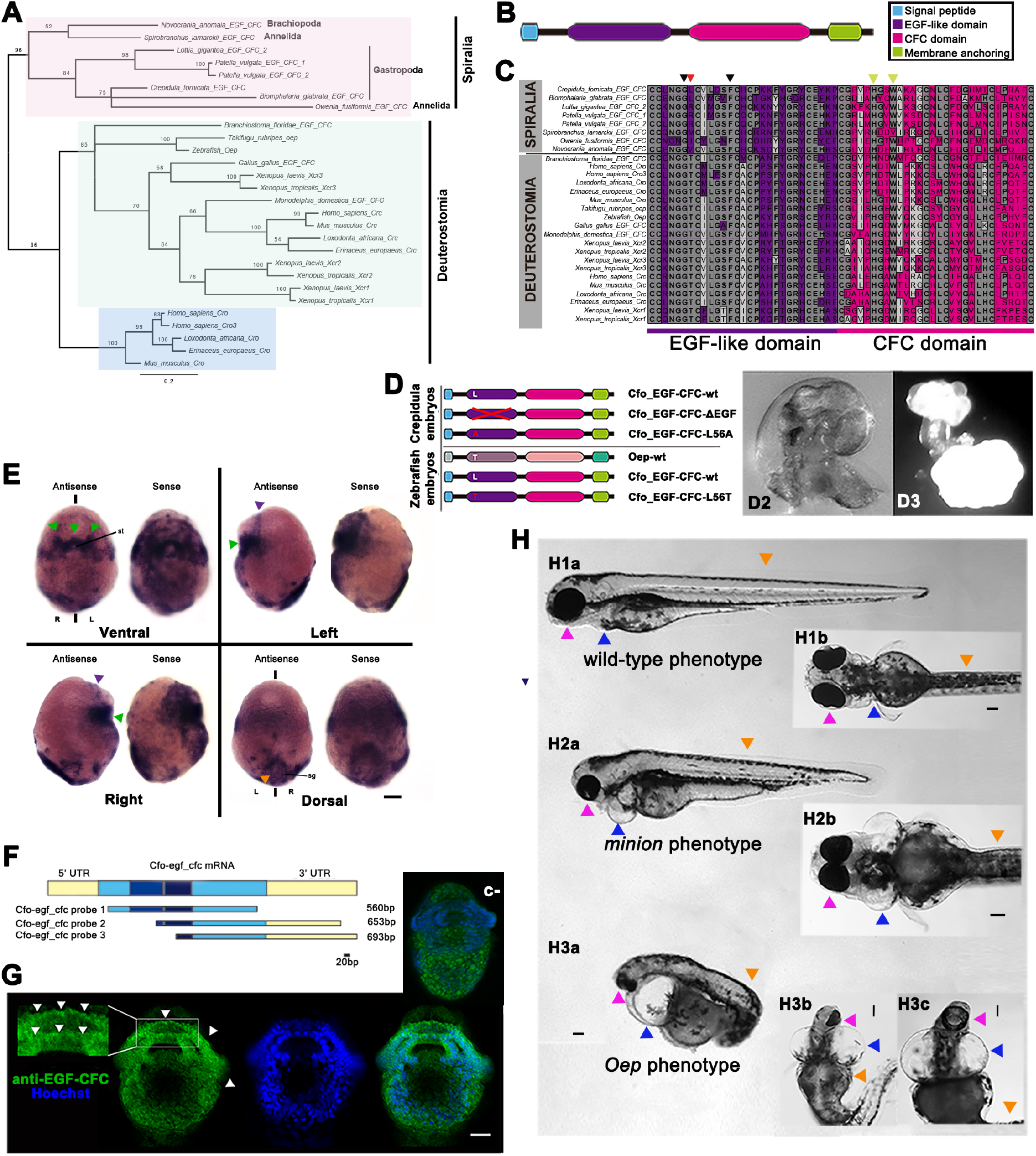
Cfo-EGF_CFC is not involved in the establishment of LRA in *C. fornicata*, but a single substitution is able to the rescue of Zebrafish Oep mutants. A) EFG-CFC proteins inter-relationships. Bayesian phylogenetic tree of EFG-CFC inferred proteins from spiralian species, along with previously described sequences of other bilaterians. The WAG model (Whelan & Goldman, 2001) was selected as the best-fit model of protein evolution using ProtTest (Abascal *et al*., 2005). Tree shown is the result of Bayesian analysis, run for 1,000,000 generations, and analyses run until the s.d. of split frequencies was below 0.01, with the first 25% of sampled trees discarded as ‘burn-in’. Posterior probabilities can be seen at each branch. Scale bar represents substitutions per site at given distances. Rooted using midpoint rooting. Shading shows major protein lineages. B) Generic structure of EGF-CFC protein. C) Alignment of the EGF-like and CFC domains of several ortholog proteins for Cripto, Cryptic and EGF-CFC in different bilaterians, from both Spiralia and Deuterostomia clades. Grey indicates conservation for one given position. Arrowheads indicate the relevant positions in the ligand-receptor-coreceptor binding; Gly and Phe (purple arrow) are conserved while Thr, described as necessary for Nodal signaling, is only present in deuterostomes and is variable in spiralians (red arrow); His and Trp (yellow arrow) are conserved in all orthologs with only one exception for *Spirobrachus*. D) mRNA constructs injected in *Crepidula* and Zebrafish embryos. D2) Left side of a wild-type *C. fornicata* juvenile; D3) Radialized individual resulting from EGF-CFC LOF, after using the construct lacking the EGF-like domain. E) Localization of *egf-cfc* transcripts by whole mount *in situ* hybridization in a *C. fornicata* preveliger larva. The left embryo on each view was incubated with the antisense probe, and the right one corresponds to the sense probe. Green arrowheads point to the symmetrical signal at the stomodeum; purple arrowhead, symmetrical anterior signals; orange, apparent symmetrical shell gland signals. st, stomodeum; sg, shell gland; L, left; R, right; bilateral axis is represented by the vertical bars. Scale corresponds to 30 μm. F) *egf-cfc* probe cloning designs. Regions in yellow represent the 5’ and 3’UTRs, and blue corresponds to the coding region. Note that the two darkest fragments correspond to the EGF-like and CFC domains. Probe 1 contains most part of the coding region, while probes 2 and 3 contain different lengths of coding region and 3’UTR. The bar indicates the scale. G) Ventral view of an immunofluorescent staining for EGF-CFC on a *C. fornicata* preveliger larva. Green channel shows the staining from a specific antibody designed against this protein, which signal appears to be symmetrical and consistent with the result of whole mount *in situ* hybridization (fig. 3E). Magnification of the stomodeum reveals this protein in the cell membranes (white arrows). The blue channel shows nuclei after Hoechst staining; followed by the merge of both channels. An equivalent larva is provided from the negative control reaction (c-); note the lack of staining at the stomodeum. Scale corresponds to 30 μm. H) Zebrafish individuals resulting from the rescue experiment on *oep* mutants, after injecting zygotes with substitution L56T on the *egf-cfc* construct. H1a) Left and (H1b) ventral views of a wild-type phenotype; H2a) Left and (H2b) ventral views of a *minion* phenotype; H3a) Left and (H3b-c) ventral views of a classic oep phenotype. Arrowheads point to the structures affected the phenotype: Pink, lens and head; blue, heart; orange, anterior-posterior axis. Scale bars correspond to 100 μm.

### *Cfo-egf_cfc* is active during development in *C. fornicata*, including the blastopore

We designed riboprobes containing the entire coding region of *Cfo-egf_cfc* and performed WISH in *C. fornicata* embryos (fig. 3F). *Cfo-egf_cfc* transcripts appear to be maternally provisioned and do not present any asymmetrical expression during development. During early development, expression becomes more intense and is restricted mainly to the region of the blastopore (data not shown). During pre-veliger stages, expression becomes very intense and is restricted to the stomodeum and part of the velum in a symmetrical pattern (fig. 3E). Although a discrete signal is also observed in cells on both left and right sides in the posterior region and in the area of the shell gland (fig. 3E), the variability between embryos did not allow us to establish a significant pattern. On the other hand, a sense probe for *Cfo-egf_cfc* also revealed expression patterns that showed little difference in localization when compared to those above (fig. 3E). This surprising result could be an artifact; however, several probes designed containing different regions of the mRNA (fig. 3F), including the 3’UTR domain, showed the same pattern. One possible explanation could be the existence of antisense RNAs, such as Natural Antisense Transcripts (NATs), that may be regulating the *Cfo-egf_cfc* gene product, and that may be encoded by the sense DNA strand (Su *et al*., 2010; Sun *et al*., 2017; Piatek *et al*., 2017). This hypothesis should be tested in future experiments.

To further analyze *Cfo-egf_cfc* expression, we designed a specific polyclonal antibody in this species to investigate specific areas of translation. Immunofluorescence experiments revealed that while control reactions without the primary antibody do not show any signal (fig. 3G c-), the pre-veliger larvae present a strong symmetrical signal located only in the membranes of the cells around the blastopore and the velar rudiment (fig. 3G). This spatial distribution is consistent with that observed in WISH results (fig. 3E); thus, confirming this data.

### Cfo-EGF_CFC is not involved in the establishment of LRA in *C. fornicata*

To investigate whether the ECF-CFC coreceptor is involved in LRA asymmetry outside Deuterostomia, we designed two approaches to interfere with its function in *C. fornicata* (fig. 3D; table 1). First, we explored gain of function (GOF) through overexpression of Cfo-EGF_CFC-wt in *C. fornicata* zygotes. No phenotypes different from the controls were observed with this approach (table 1). Second, we explored the LOF through two designs for defective forms, after studying the residues involved in the Nodal pathway. As indicated before, the EGF-like domain is known to be involved in binding the ligand. Initially, we designed a form of Cfo-EGF_CFC devoid of the EGF-like domain (Cfo-EGF_CFC-ΔEGF) (fig. 3D; table 1). This experiment resulted in an increased mortality rate and those that survived displayed a radialized phenotype, characterized by the absence of dorso-ventral and left-right axes (fig. 3D3). This result differs from those of Nodal GOF and LOF, suggesting a possible interference with other pathways that this coreceptor is known to regulate. The resulting radialized embryos, which did not undergo gastrulation, had no axes and diminished cell differentiation, are reminiscent of the phenotype obtained when the MAPK pathway is inhibited in *C. fornicata* (Henry & Perry, 2008). This, together with the fact that EGF-CFC activates the MAPK pathway in vertebrates (Kannan *et al*., 1997), suggests that the MAPK pathway may be regulated by Cfo-EGF_CFC in *C. fornicata*. However, more experiments should be performed to test this hypothesis.

Since the EGF-CFC coreceptor could be binding different ligands and it is known that specific residues are responsible for each of them, we focused on finding the orthologous amino acid to that located at position 88 (T88) in humans, which is described as critical for Nodal interaction in vertebrates. As described above, this position corresponds to a leucine in *C. fornicata* (L56) (fig. 3C). Hence, we first had to determine whether this amino acid position was as crucial for Cfo-EGF_CFC function in gastropods as T88 was in vertebrates. Finding a leucine in this relevant position made us suspect that the coreceptor may not be able to interact with the ligand Nodal, but we also considered the probability of having other interactions involving this change. Therefore, we generated a mutation at L56 by substituting the leucine with alanine L56A (Cfo-EGF_CFC-L56A) and injected this mRNA in *C. fornicata* zygotes (table 1). No phenotype different from a normal wild-type was observed in the larvae. Although we conclude that this amino acid has no impact in LRA in *C. fornicata*, we could not speculate whether this change to leucine has any real implications on Nodal pathway signaling and LRA.

### L56T substitution promotes a gain of function in EGF-CFC that can rescue a deuterostome mutant line for its ortholog

After the results described above with Cfo-EGF_CFC-L56A, we reasoned that if L56 is a true equivalent to position T88 in deuterostomes, the L56A mutant form should be completely unable to function. The best way to test this hypothesis would be by attempting to rescue an *egf-cfc* loss-of-function mutant by overexpressing the wild-type construct of *C. fornicata*. Since we do not have such a mutant in *C. fornicata*, we turned to zebrafish (fig. 3D). An extensively characterized mutant line in the *egf-cfc* gene exists in zebrafish (*oep*). Matings of heterozygous parental individuals display a typical Mendelian segregation (table 3), in which 25% of the embryos show an *oep* phenotype (*oep*-/-) (fig. 3H) and 75% of the embryos present a wild-type phenotype (*oep* +/-; *oep*+/+) (fig. 3H1a-b). Oep mutants display cyclopia (presence of just one eye due to an undivided forebrain and orbital cavity), anterior-posterior axis deviation with shortening, cardiac hypertrophy and inversion of the heart (Schier *et al*., 1997). We first confirmed that injection of a wild-type form of zebrafish *oep* mRNA can rescue the mutant phenotype, as previously described. We synthesized the mRNA for *D. rerio oep*, and injected it into zygotes obtained from heterozygous matings (table 3). The results showed an increased number of wild-type phenotypes (89%) in comparison to the expected scoring under normal conditions (75.9%) (table 3). Next, we used the two constructs designed for *C. fornicata* Cfo-EGF_CFC-wt and Cfo-EGF_CFC-L56A. Additionally, we created a third mutation, leucine to threonine (Cfo-EGF_CFC-L56T), that attempts to reproduce the form found in deuterostomes. The injection results in zygotes from the same Zebrafish mating are shown in fig. 3H and table 3. Our findings showed that Cfo-EGF_CFC-wt could not rescue the *oep* phenotype (21.6% mutants) (table 3), nor the presumptive defective form Cfo-EGF_CFC-L56A (data not shown). However, Cfo-EGF_CFC-L56T rescued the phenotype (10.3% mutants) in a proportion similar to wild-type zebrafish *oep* (11% mutants) (table 3). In some of the cases, a partial rescue that represents an intermediate phenotype was detected and referred to as the “Minion” phenotype, which displays a significant rescue of bilaterality (fig. 3H2a-b). The occurrence of the Minion phenotype strengthens our hypothesis of a recovery of function in the Cfo-EGF_CFC-L56T injections, since this showed a noticeable reduction of the cardiac hypertrophy, an apparent complete recovery of the anterior-posterior axis, nearly complete development of the head and the generation of two eyes with partial cyclopia (closer set compared to those in the wild-type embryos) (fig. 3H). Overall, these results suggest that even though an EGF_CFC ortholog is present in *C. fornicata*, the product of this gene is unable to modulate the Nodal pathway due to an inability to bind to Nodal.

**Table 3.** Results obtained after mRNA injection of different *egf-cfc* constructs, in zygotes of the *D. rerio* oep mutant line.

|  | mRNA & dose | Total embryos | Observed phenotype |  |  |
| --- | --- | --- | --- | --- | --- |
|  |  |  | Wild-type | Oep | Minion |
| Control experiment (F1 for oep +/- cross) | - | 54 | 41 (75.9%) | 13 (24.1%) | 0 |
| Rescue with *Zf_oep* gene | Oep-wt | 73 | 65 (89%) | 8 (11%) | 0 |
| Rescue with *Crepidula* wt | Cfo_egf-cfc_wt | 97 | 76 (78.4%) | 21 (21.6%) | 0 |
| Rescue with *Crepidula* L56T mutant | Cfo_egf-cfc_L56T | 39 | 33 (84.6%) | 4 (10.3%) | 2 (5.1%) |

## DISCUSSION

Nodal has been studied in detail for decades, and research has been extended to non-deuterostomes since its discovery in snails in 2009 (Grande & Patel, 2009). The comparative analysis in this study demonstrated conservation of the Nodal pathway and its function among the Bilateria. However, the regulatory aspects of the Nodal pathway in gastropods, as well as the effects on its perturbation remain unexplored. Our study provides the first evidence of the presence of an EGF-CFC coreceptor on non-deuterostome representatives, demonstrating that the EGF-CFC family traces back to the last common ancestor of Bilateria. Our functional analysis supports the hypothesis that EGF-CFC had an ancestral role unrelated to the Nodal pathway. We propose that a gain of function event driven by a single amino acid change was responsible for facilitating a new role for the coreceptor within the Nodal pathway.

### Nodal and EGF-CFC were present in the ancestor of Bilateria

As previously reported by Grande *et al*. (2014), Nodal is present in most bilaterians, and two paralog genes were found for *C. fornicata* (fig. 1A, fig. 4). Studies focusing on the origin and molecular evolution of the EGF-CFC coreceptors identified several members of this family in deuterostomes, which led some authors to conclude that this family originated in the deuterostome ancestor (Ravisankar *et al*., 2011). The data presented here demonstrate the existence of the EGF-CFC proteins in spiralians (figs. 3A and 3C, fig. 4) tracing this coreceptor back to the last ancestor of all bilaterians. Besides *Patella*, all the non-vertebrate organisms included in the analysis present only one ortholog. These results can be explained by two gene duplication and divergence events, one in *Patella* and a second in the ancestor of all deuterostomes. Subsequently, one of the copies in the deuterostome lineage was lost in some groups such as fish, *Gallus* or marsupials. Additionally, broader sampling in other groups of metazoans is needed to make real inferences of the evolutionary history of these proteins.

**Figure 4.**
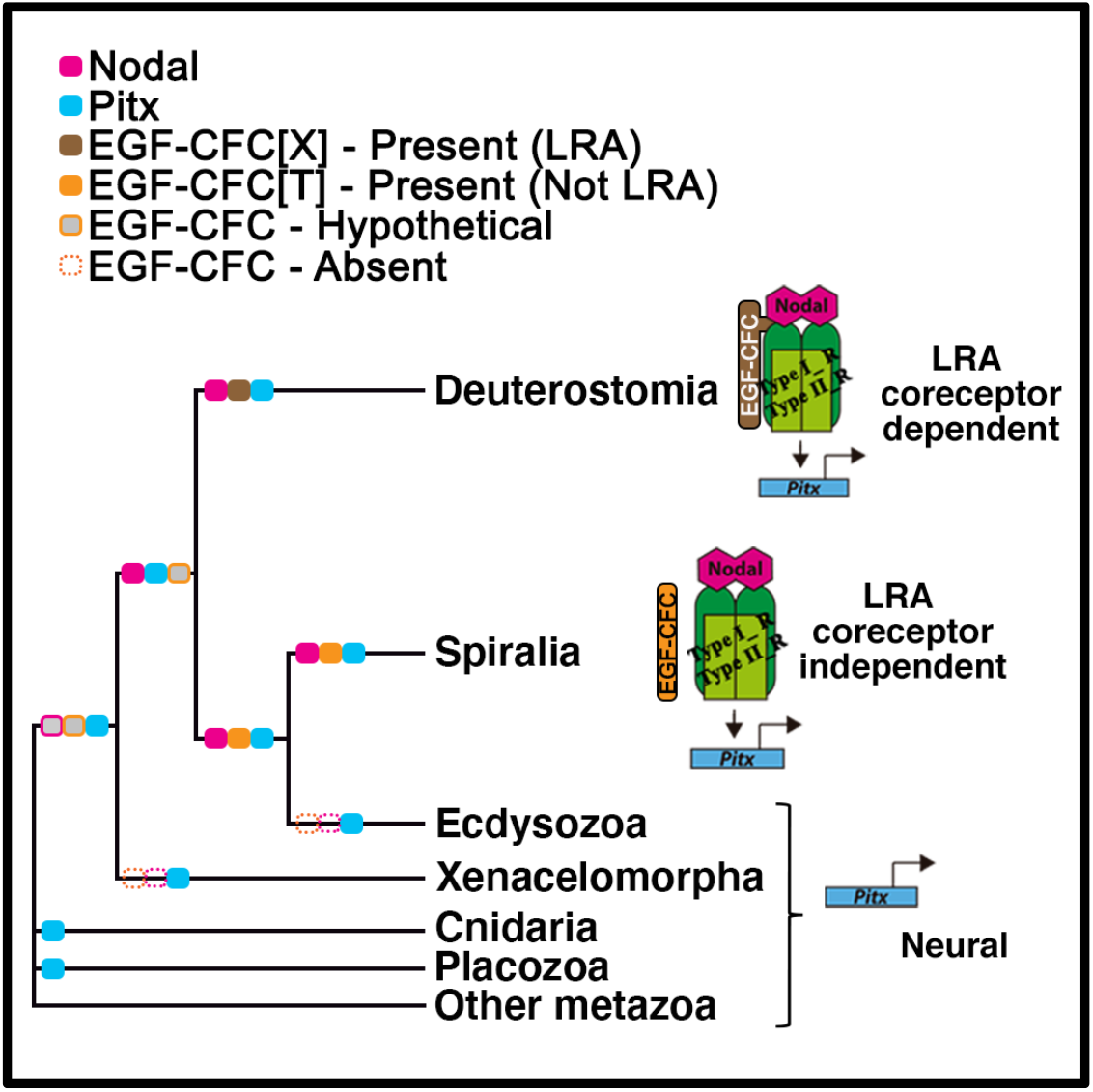
Proposed model for the evolution of EGF-CFC role in Metazoa. The orthologs for EGF-CFC in Spiralian representatives display different amino acids on this relevant position in the EGF-like domain, whereas all representatives of Deuterostomia display a Threonine. The change of amino acid in this specific position would have happened in the ancestor of Deuterostomia. This gave the coreceptor the ability to interact with the ligand, and regulate Pitx expression, thereby participating in the establishment of LRA. In Spiralia and its ancestor, the function of EGF-CFC would not be linked to LRA, but to gastrulation and axis determination. The regulation of LRA is proposed to be independent of the coreceptor for Spiralia. Pitx ancestral function would be related to neural specification, and it would remain as the only function of this gene in those clades that lack the ligand and co-receptor. The color filled boxes represent presence of the protein in a given clade. The grey filled boxes represent inferred hypothetical presence of the protein. The dotted line boxes represent the absence of an ortholog. X, any amino acid; T, Threonine; LRA, Left-right asymmetry.

### Nodal and EGF-CFC are expressed during development in *C. fornicata*

As observed in previous work in *P. vulgata* (Kenny, 2014), in *C. fornicata* only one *nodal* paralog displays asymmetrical expression from early stages, and remains asymmetrically active during development; whereas the second paralog does not (figs. 1C and 1D). Gene duplication events can lead to paralog sub-functionalization and the emergence of new functions. Further comparative analysis will provide interesting data in relation to these events. Expression domains of *nodal* in the Bilateria seem to be highly divergent (Hamada *et al*., 2002; Grande & Patel, 2009; Martín-Durán *et al*., 2016). The early asymmetrical expression of *Cfo-nodalB* is consistent for non-deuterostome clades (Grande & Patel, 2009; Martín-Durán *et al*., 2016), but differs from the asymmetrical establishment of Nodal deuterostomes. Indeed, in deuterostomes nodal expression is initially symmetrical and transitions into an asymmetrical pattern through several regulatory mechanisms (Duboc *et al*., 2005; Ermakov, 2013). It remains unclear what mechanisms activate *nodal* expression directly in an asymmetrical way in spiralians, or whether those activators are shared with deuterostomes, and if so, how they evolved and diverged. Consequently, clear differences appear to exist related to the regulation of *nodal* expression among different bilaterians. On the other hand, we considered that *Cfo-nodalB* could be related to LRA given its gene expression pattern during development (fig. 1C).

The Nodal target *Pitx* is also asymmetrically expressed following *nodal* activation (Grande & Patel, 2009; Hamada *et al*., 2002). Some domains of *Pitx* expression patterns are symmetric in *Crepidula* (fig. 1E), as is the case in other gastropods and vertebrates, suggesting the presence of independent enhancer areas that regulate this expression (Christiaen *et al*., 2005; Grande & Patel, 2009). *Pitx* is the only target described so far in gastropods. Identifying additional targets will provide further insight about body plan evolution.

Our analysis revealed that *egf-cfc* is symmetrically expressed around the blastopore lip in *C. fornicata* (fig. 3E). This structure is a key signaling center during gastrulation in gastropods (Van den Biggelaar *et al*., 2002). Hence, these results show that the *egf-cfc* gene may have a relevant role during gastrulation in gastropods, as is the case in vertebrates (Ding *et al*., 1998; Chu *et al*., 2005). Despite our efforts and region/condition modifications, both the antisense and sense probes showed the same restricted and recurring pattern (figs. 3E and 3F). These results suggest that there may be an antisense and a sense transcript, both active during *Crepidula* development. Natural antisense transcripts, NATs, are RNA molecules complementary to the sense mRNA, endogenously found and active in eukaryotes and “prokaryotes”. NATs have been found to influence embryonic development in some vertebrate species (reviewed in Su *et al*., 2010; Piatek *et al*., 2017; Sun *et al*., 2017). Up to 72% of mRNAs have an antisense RNA in mouse and human transcriptomes (Beiter *et al*., 2009). It is plausible that the *egf-cfc* sense transcript in *C. fornicata* may be regulated by an antisense transcript. Nevertheless, more experiments would be necessary to properly validate this hypothesis. To overcome the limitations of our expression analysis, a specific antibody was designed for this work. Immunostainings are consistent with the WISH experiment, given the presence of signal in the membranes of the cells surrounding the blastopore lip in larvae. This signal spreads to the anterior velar rudiment region and along both sides in a symmetrical manner (fig. 3G), and supports the potential relevance of EGF-CFC during gastrulation in gastropods.

### The role of Nodal and EGF-CFC in establishment of LRA

GOF and LOF experiments confirm that Nodal is involved in LRA in *C. fornicata*, as previously reported for deuterostomes (Hamada, *et al*., 2002) and other non-deuterostomes (Grande & Patel, 2009). Ectopic activation of the Nodal pathway leads to loss of asymmetric coiling shell growth, and the characteristic twist of the digestive tract (fig. 2B; table 1). This result is consistent with previous reports of loss of bilateral specialization by side identity disruption in vertebrates (Hamada *et al*., 2002; Duboc *et al*., 2005), and experiments on differential growth rate of shell asymmetrically produced by the shell gland cells (Grande & Patel, 2009; Kurita & Wada, 2011). On the other hand, the lack of phenotype for the LOF construct missing the cleavage site questions the functionality of the design (table 1). LOF by RHAct-A confirmed the importance of this pathway for LRA, since *Crepidula* embryos at 30 hpf treated for two days led to 80% of symmetric juveniles (fig. 2A, 2C; table 2). Moreover, these embryos did not show any asymmetrical expression of *Pitx* compared to controls (fig. 2D). These data indicate that the RHAct-A treatment is an effective approach to block the Nodal pathway in *Crepidula*, and leads to LRA loss (Supp. material S1). Interestingly, the symmetric expression domains of *Pitx* remained comparable to those detected in untreated embryos (fig. 1C), and are consistent with LOF experiments previously reported (Grande & Patel, 2009). We hypothesize that a different combination of enhancers may be controlling the expression of *Pitx* in the region symmetrically expressing the gene, or even cross-talk with another pathway could regulate its expression in these regions unaffected by RHAct-A (i.e., Wnt, Notch (Luo *et al*., 2017)). This Pitx-positive region could be related to the possible ancestral role of Pitx, presumably related to neural specification. The appearance of Nodal could have promoted the co-option of the *Pitx* gene for a new function, now involving it in LRA (Grande *et al*., 2014) (fig. 4).

Our results suggest that EGF-CFC, described as a master regulator of the pathway in deuterostomes (Wechselberger *et al*., 2005; Kelber *et al*., 2008), does not have a role in LRA, nor does it regulate Nodal signaling in *C. fornicata* (fig. 4). However, the data regarding its expression and protein location together with LOF experiments performed through EGF-CFC_Cfo_ΔEGF (fig. 3D; table 1), suggests a conserved role in gastrulation and body axis formation. The key amino acid for Nodal pathways is leucine in *C. fornicata* (fig. 3D). However, the mutation at this position did not produce any phenotype (table 1), suggesting that the ancestral regulation of Nodal signaling was independent of the coreceptor, and an EGF-FCF-dependent pathway would have evolved only in deuterostomes (fig. 4). Indeed, this is supported by previous evidence that Nodal has Cripto-independent activity in vertebrates, and the proposal of the existence of two pathways for this morphogen (a Cripto-dependent and a Cripto-independent pathway) (Brennan *et al*., 2001; Liguori *et al*., 2008).

### The connection between Nodal and EGF-CFC emerges in the ancestor of deuterostomes

We explored the ability of the *C. fornicata* ortholog to rescue the zebrafish mutant line for EGF-CFC, *oep* (fig. 3D). We reasoned that this approach would allow us to determine the conservation in the function of EGF-CFC across phyla. Overexpression of the wild-type form of *C. fornicata* EGF-CFC does not rescue the *oep* phenotype. Remarkably, changing a single amino acid (L56T) in *C. fornicata* EGF-CFC is necessary and sufficient to restore EGF-CFC activity and rescue *oep* mutants (fig. 3H; table 3). We propose that, although Nodal and EGF-CFC were present in the last ancestor of Bilateria, their functional connection occurred in the deuterostome stem lineage. Prior to that, EGF-CFC may have had an ancestral role unrelated to Nodal, but important for gastrulation and body axis formation. The presence of threonine appears to represent a change that occurred in the deuterostome common ancestor, and correlates with a novel function for this vital coreceptor and a new GRN for the Nodal ligand.

## MATERIAL AND METHODS

- Full version available after acceptance for publication.

### Experimental model and subject details

All animal experiments were performed in accordance with the governmental and institutional guidelines. *C. fornicata*-Adults of *C. fornicata* were harvested from local waters near Woods Hole, MA by the MRC at the MBL. Embryos were collected and reared as previously described (Henry *et al*., 2010; Perry *et al*., 2015; see also https://www.life.illinois.edu/henry/tools.html). Embryos and larvae were fixed as described in Perry *et al*. (2015). *D. rerio -* AB and tupl wild-type zebrafish strains, and the *oep*^*tz57*^ mutant line were maintained and bred according to standard procedures. Embryos were harvested and grown in system water at 28ºC in a humid incubator. All experiments conform to the guidelines from the European Community Directive and the British and Spanish legislation for the experimental use of animals.

### Sequence analysis and orthology establishment

Available after acceptance for publication.

### Cloning, probe synthesis, whole-mount *in situ* hybridization

High quality total RNA was extracted and purified as previously described (Kenny *et al*., 2016). cDNA was generated by using SMARTer PCR cDNA Synthesis Kit (Clontech). Gene-specific primers were designed for each gene of interest (Supp. material S3). PCR products were cloned into the pGEM-T Easy vector (Promega, Madison, WI). Linearized template DNA (amplified from plasmid with T7/SP6 primers) was used to synthesize RNA probes with either T7 or SP6 RNA polymerase (Life Technologies), and DIG-labeling mix (Roche, Indianapolis, IN). WISH protocol was performed, as previously described (Perry *et al*., 2015).

### mRNA templates, transcription, microinjection

The mRNA was prepared by PCR amplification of the construct, with SP6 and T3 primers and Phusion High Fidelity DNA polymerase (New England, BioLabs). The purified product was used as a template for the transcription reaction using the mMessage mMachine SP6 RNA transcription kit (AM1340, Ambion, Austin, TX), as previously described (Henry *et al*., 2010). The same protocol was followed for *oep* mRNA, amplified from cDNA generated from 24 hours post-fertilization *D. rerio* embryos. Fertilized eggs of *C. fornicata* and *D. rerio* were pressure microinjected. The device used for the experiments on *C. fornicata* is described in Truchado-Garcia *et al*., 2018 (Truchado-Garcia M, Harland RM, Abrams MJ, unpublished method, https://www.biorxiv.org/content/10.1101/376657v1.full), and a semi-dry technique (pretri-dish and a glass slide) was used to hold zebrafish embryos.

### Antibody production and antibody staining

To generate Cfo_EGF-CFC-specific antibodies, a peptide of this protein was synthetized and inoculated in rabbits, by 21st Century Biochemicals, Inc. Resulting polyclonal antibodies were subsequently purified by the same company. Whole specimens were permeabilized on PBT (0.2% X-Triton100 in 1xPBS), and blocked in 2% BSA in PBT. Sample was incubated with the primary antibody (anti-Cfo_EGF-CFC 1:10), overnight at 4C. Several washes in PBT were performed prior incubation with the secondary antibody (goat anti-IgG rabbit 488, Molecular Probes, A-11008), at room temperature for 2 hours. Nuclei were stained with Hoechst 1:1000 (Molecular Probes, H3570) for 30 minutes in PBT, at room temperature and darkness. Last, specimens were washed several times in PBT and final in 1xPBS. Samples were stored in 80% glycerol-PBS for imaging.

### Recombinant human Activin treatment in *C. fornicata*

*C. fornicata* embryos at 3 different stages (two-cell stage (7-8 hours post-fertilization; hpf), morula (32 hpf), early gastrula (48 hpf; 45% epiboly) and late blastula (61hpf; 65% epiboly)) were treated with 5 or 10 µM human recombinant Activin-A protein, or AFSW (Artificial Filtered Sea Water) with 0.1% DMSO, as a control. Embryos were cultured as previously described (Henry *et al*., 2010) for 48 hours, after which point the protein was washed out by replacement with 100% AFSW, repeated twice.

### Microscopy

Slides were prepared as previously described in Perry *et al*. (2015). Brightfield imaging was carried out on a Zeiss Axioskop2 plus (Carl Zeiss, Inc., Munich, Germany). Images were captured with a CoolSnap FX color camera (Roper Scientific) running Metavue 7.10 software (Universal Imaging). Multifocal stacks were processed using Helicon Focus stacking software 6.7.1 (Helicon Soft Ltd., Kharkov, Ukraine). Confocal imaging was carried out on a LSM710 AxioImager M2 (Zeiss), running Zen 2010 software (Zeiss). Snail larvae were anesthetized with magnesium chloride, as previously described (Perry *et al*., 2015) and imaged with a DFC350Fx camera docked to a M205FA scope (Leica Microsystems). *D. rerio* embryos were anesthetized in 0.01% tricaine. ImageJ v2.0.0-rc-56/1.51h and Adobe Photoshop CS5 extended v12.1 x64 were used for processing the images.

## Acknowledgements

We thank the Marine Biological Laboratory and specially the former directors of the Embryology course R. Behringer and A. Sánchez Alvarado, for supporting this research, J.M. Martín-Duran for helping with the searches for EGF-CFC orthologs on the RNA-seq data of some bilaterians,

M. J. Abrams and R. M. Harland for helping with edition and discussion. This work was supported by the Spanish Ministry of Science, Innovation and Universities (MICINN) (grant CGL201-29916 to CG; predoctoral fellowship BES-2012-15 052214, and short-time appointment fellowships EEBB-1-14-08959, EEBB-1-15-16 09637, EEBB-1-16-11411 to MT-G to work with JQH and KJP). CG was for a portion of the time spent on this project a “Ramon y Cajal” postdoctoral fellow supported by the Spanish Ministry of Economy and Competitiviness (MEC) and the Universidad Autonoma de Madrid (UAM) (Spain).

## SUPPLEMENTAL MATERIAL 1

**S1.**
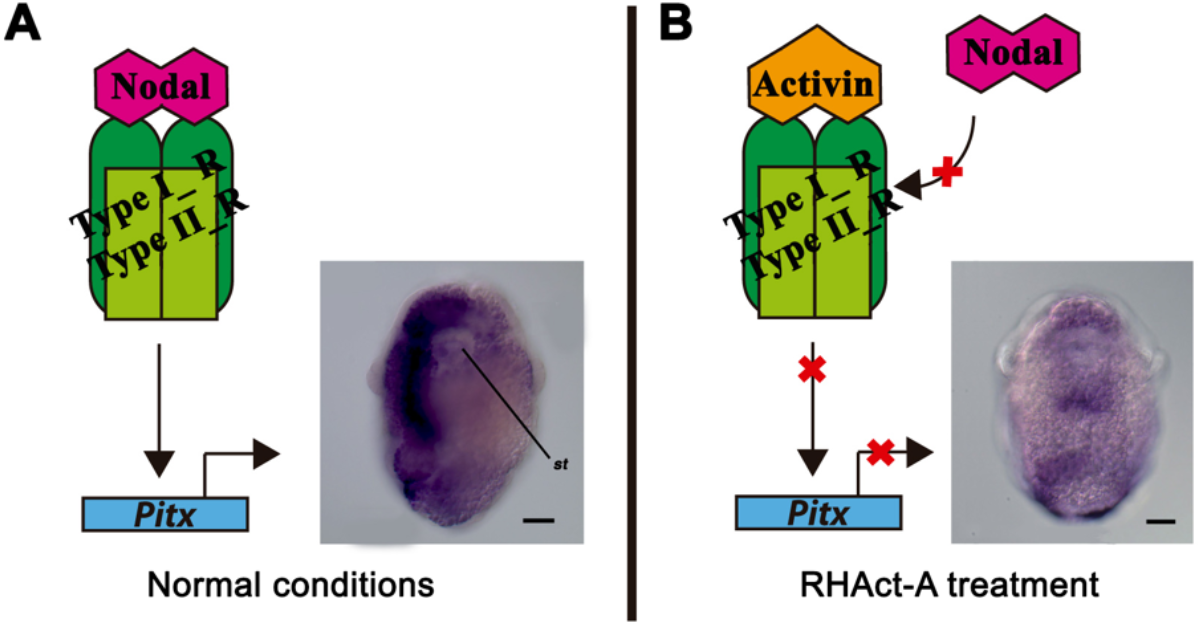
The addition of RHAct-A leads to the inactivation of the Nodal pathway, presumably by competition with the ligand NodalB for the receptors. A) Transcription of *Pitx* is activated by the signaling cascades transduced the NodalB binding to the receptor cluster. On preveliger larvae, *Pitx* is symmetrically expressed on the anterior region and asymmetrical on the right side (fig. 1E). B) RHAct-A presumably prevents NodalB from binding the receptor cluster, due to direct competition. Subsequently, no *Pitx* expression is promoted, as showed by *Pitx* whole mount *in situ* hybridization on treated embryos; though such silencing only occurs in the asymmetrical territories, while its symmetrical activation signals remain unaffected. Scale corresponds to 30 μm.

